# Kif18A promotes Hec1 dephosphorylation to coordinate chromosome alignment with kinetochore microtubule attachment

**DOI:** 10.1101/304147

**Authors:** Haein Kim, Jason Stumpff

**Affiliations:** Department of Molecular Physiology and Biophysics, University of Vermont, Burlington, VT 05405, United States; Lead Contact

## Abstract

Mitotic chromosomes are spatially confined at the spindle equator just prior to chromosome segregation through a process called chromosome alignment. Alignment requires temporal coordination of kinetochore microtubule attachment and dynamics. However, the molecular mechanisms that couple these activities are not understood. Kif18A (kinesin-8) suppresses the dynamics of kinetochore microtubules to promote chromosome alignment during metaphase. Loss of Kif18A function in HeLa and primordial germ cells leads to alignment defects accompanied by a spindle assembly checkpoint (SAC)-dependent mitotic arrest, suggesting the motor also plays a role in regulating kinetochore-microtubule attachments. We show here that Kif18A increases attachment by promoting dephosphorylation of the kinetochore protein Hec1, which provides the primary linkage between kinetochores and microtubules. This function requires a direct interaction between the Kif18A C-terminus and protein phosphatase-1 (PP1). However, the Kif18A-PP1 interaction is not required for chromosome alignment, indicating that regulation of kinetochore microtubule dynamics and attachments are separable Kif18A functions. Mitotic arrest in Kif18A-depleted cells is rescued by expression of a Hec1 variant that mimics a low-phosphorylation state, indicating that Kif18A-dependent Hec1 dephosphorylation is a key step for silencing the checkpoint and promoting mitotic progression. Our data support a model in which Kif18A provides positive feedback for kinetochore microtubule attachment by directly recruiting PP1 to dephosphorylate Hec1. We propose that this function works synergistically with Kif18A’s direct control of kinetochore microtubule dynamics to temporally coordinate chromosome alignment and attachment.

## Introduction

Kinetochores, which form at the centromeric region of mitotic chromosomes, function to tether chromosomes to a subset of mitotic spindle microtubules known as kinetochore microtubules or K fibers. Proper regulation of K fiber dynamic instability and attachment is required for chromosome alignment at metaphase and the segregation of sister chromatids during anaphase [1–4]. K fiber attachments are stabilized as mitotic chromosomes align [5–7], but the mechanisms that temporally coordinate attachment and alignment are not understood.

End-on attachments between K fibers and kinetochores are primarily dependent on Hec1, a component of the NDC80 complex. Hec1 directly associates with microtubules through an electrostatic interaction that depends on its positively charged N-terminus [8, 9]. The Hec1 N-terminus contains nine Aurora B phosphorylation sites, which are phosphorylated in early mitosis [10–13]. Hec1 phosphorylation reduces the interaction between the kinetochore and the negatively charged microtubule surface, favoring K fiber detachment [2, 9, 14–16]. Hec1 is dephosphorylated as cells progress through mitosis, with the lowest levels occurring during late metaphase and anaphase [14]. These data indicate the affinity of kinetochores for microtubules increases as chromosomes align. Recruitment of protein phosphatase 1 (PP1) to kinetochores by proteins such as KNL1 [17, 18], and the SKA complex [19] has been shown to antagonize Aurora B activity, however, the specific mechanism responsible for Hec1 dephosphorylation remains unclear.

Hec1 affinity for microtubules must be dynamically regulated, not only for proper biorientation of chromosomes, but also for normal chromosome alignment. Cells microinjected with antibodies that block Hec1 phosphorylation show hyperstable K fiber attachments, accompanied by alignment defects and decreased chromosome oscillations [2]. Similarly, cells expressing a Hec1 variant that prevents phosphorylation of the N-terminus are unable to align their chromosomes due to reduced K fiber dynamics [16, 20]. Conversely, in cells expressing a phosphomimetic Hec1 variant, with the 9 Aurora B sites in the N-terminus mutated to acid residues, kinetochores are uncoupled from K fibers and chromosomes fail to align [16, 20]. Furthermore, introduction of a single phosphomimetic residue to the non-phosphorylatable Hec1 variant (Hec1-1D) is sufficient to restore normal chromosome movement but not chromosome alignment [19, 20], suggesting that proper timing of Hec1 dephosphorylation is required for chromosome confinement to the metaphase plate. Consistent with this, PP1 accumulation at kinetochores increases as cells progress towards metaphase [21], coinciding with a decrease in phospho-Hec1 levels [14]. However, the mechanisms underlying temporal changes in Hec1 affinity and PP1 recruitment are not understood.

Kif18A (kinesin-8) is essential for chromosome alignment during metaphase and directly binds PP1 [4, 22–24]. This plus-end directed motor accumulates at the ends of K fibers and attenuates their dynamics as cells progress from prometaphase to metaphase. Reduced K fiber dynamics confines chromosome movements and promotes metaphase plate formation [4]. In addition, Kif18A depletion in HeLa cells results in a spindle assembly checkpoint (SAC)-dependent metaphase arrest [4, 25, 26], which suggests a defect in kinetochore microtubule attachment. Similarly, loss of Kif18A function leads to a SAC-dependent mitotic arrest in primordial germ cells during murine embryogenesis [27]. Whether Kif18A has a direct or indirect role in promoting kinetochore microtubule attachments in these cells types is unclear. However, the fission yeast kinesin-8s Klp5 and Klp6 are required for SAC silencing through an unknown function that relies on direct interaction between the C-termini of the motors and PP1 [24]. Human Kif18A contains the conserved R/VxVxF/W PP1 binding motif found in Klp5 and Klp6 and directly binds PP1α/γ [22, 23]. The Kif18A-PP1 interaction has been proposed to antagonize phospho-inhibition of Kif18A by Cdk1, but a role for this interaction in promoting kinetochore microtubule attachments and the metaphase-to-anaphase transition has not been thoroughly explored.

In this study, we used a combination of quantitative immunofluorescence and livecell imaging techniques to investigate the molecular basis of Kif18A’s role in coordinating chromosome alignment with the metaphase-to-anaphase transition. We report that Kif18A-PP1 binding promotes Hec1 dephosphorylation and progression through mitosis. This mechanism, in combination with Kif18A’s direct regulation of chromosome movement, coordinates chromosome alignment with the stabilization of K fiber attachments, promoting timely transition from metaphase-to-anaphase.

## RESULTS

### Kif18A depletion increases Hec1 phosphorylation during metaphase

Observations that Kif18A is required for normal mitotic progression in HeLa and primordial germ cells suggest that the motor has a role in promoting or stabilizing kinetochore-microtubule attachments [26, 28]. To determine if Kif18A affects the phosphoregulation of Hec1, which is progressively dephosphorylated to increase the affinity of kinetochores for microtubules as chromosomes align, we analyzed Hec1 phosphorylation by immunofluorescence in HeLa cells treated with scrambled control (control KD) and Kif18A specific siRNAs (Kif18A KD), which we have extensively validated [4, 29, 30]. Previous work has shown that the signals from Hec1 antibodies against phosphorylated Ser8, Ser15, Ser44, and Ser55 decrease significantly between prometaphase and metaphase, with Ser44 and Ser55 showing the most dramatic changes [14]. Thus, we quantified the signal from a phospho-specific antibody against Hec1 Ser55 in metaphase arrested cells to determine if Kif18A is required for Hec1 dephosphorylation (Figure 1A). These studies revealed that Kif18A KD cells have a significantly higher level of phosphorylated Hec1 at metaphase kinetochores than control cells (Figure 1B). In contrast, total Hec1 levels were comparable between the two cell populations (Figure 1C). These data indicate that Kif18A is required to promote Hec1 dephosphorylation.

**Figure 1.**
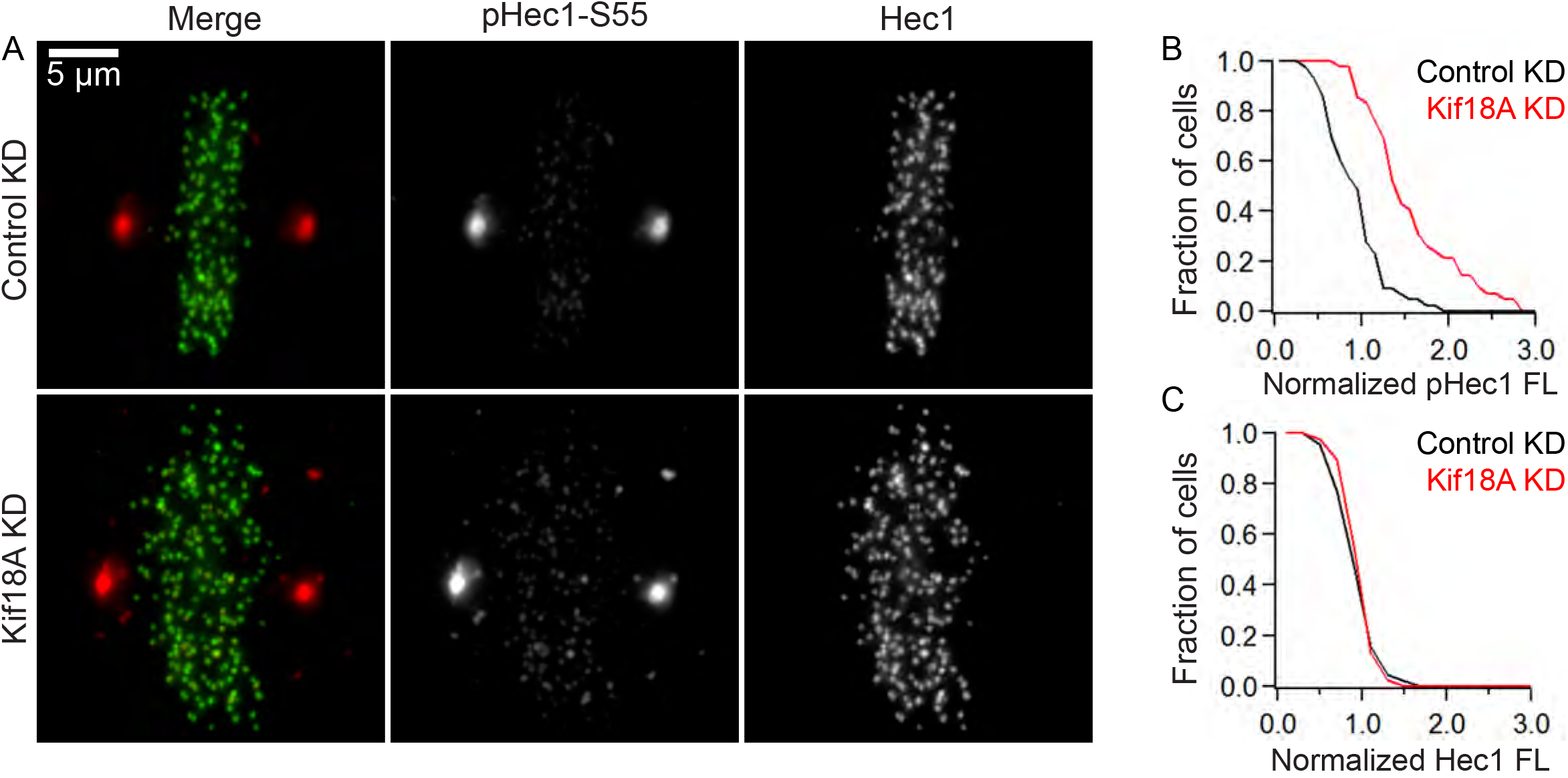
Kif18A depleted HeLa cells show a significant increase in phospho-Hec1 levels at kinetochores. **A**. Immunofluorescence microscopy images of cells labeled with total Hec1 (green) and phospho-Hec1 Ser55 (pHec1-S55, red) antibodies. Cells were treated with control (Control KD) or Kif18A siRNAs (Kif18A KD), then arrested in late metaphase with MG132 for 3 hours and fixed. **B-C**. Quantification of phospho-Ser55 (B) and total Hec1 fluorescence (C) at kinetochores in Control KD and Kif18A KD cells normalized to average Control KD levels. Data are from 3 independent experiments: n = 43 cells, 2100 kinetochores for Control KD, n = 42 cells, 2000 kinetochores for Kif18A KD, A student’s t-test was used to compare normalized fluorescence intensities (p < 0.0001, phospho-Ser55 fluorescence and p > 0.05, total Hec1 fluorescence).

To determine if Kif18A dependent changes in phospho-Hec1 levels were due to the motor’s previously identified interaction with PP1 [22, 23, 31], we quantified the effects of disrupting Kif18A-PP1 binding on Hec1 phospho-Ser55 levels. HeLa cells depleted of endogenous Kif18A were transfected with plasmids encoding GFP alone, GFP-tagged Kif18A full-length protein (GFP-Kif18AFL), or a GFP-Kif18A construct containing two point mutations within the conserved PP1 binding site (GFP-Kif18A^AVVVA^). These mutations have previously been shown to disrupt binding between kinesin-8 motors and PP1 [22–24]. Both GFP-Kif18A constructs localized to mitotic spindles and accumulated at K fiber plus-ends (Figure 2A). However, phospho-Ser55 levels were significantly lower in cells expressing GFP-Kif18AFL compared to those expressing similar levels of GFP-Kif18A^AVVVA^ (Figure 2B-C and Fig S1). Furthermore, phospho-Ser55 levels in GFP-Kif18A^AVVVA^ expressing cells were comparable to those measured in cells expressing GFP only (Figure 2B-C). These data indicate that Kif18A promotes Hec1 dephosphorylation through its interaction with PP1. This effect may not have been detected in a previous study due to utilization of a less sensitive measurement approach [23].

**Figure 2.**
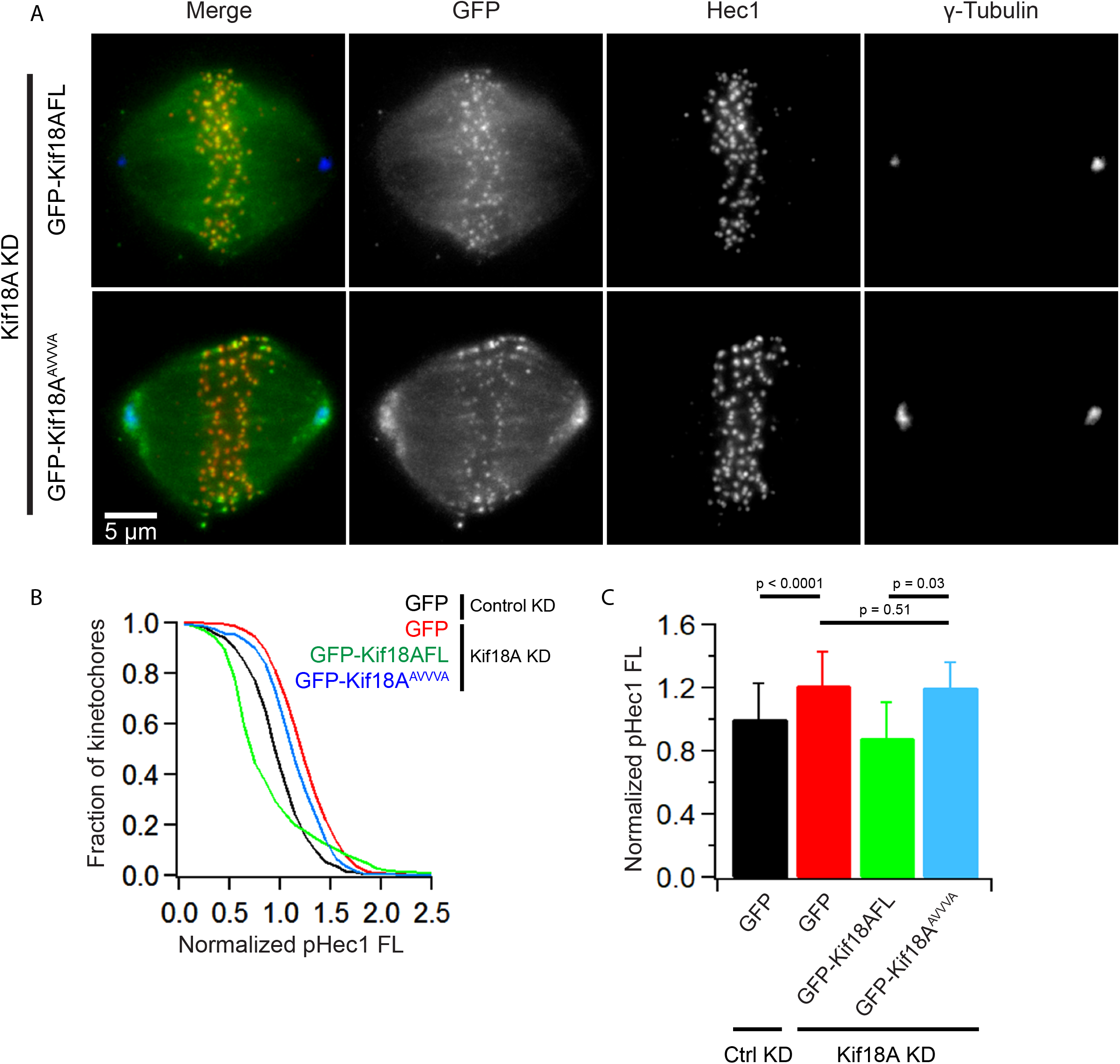
PP1 binding is required for Kif18A-dependent changes in Hec1 phosphorylation. A. Representative images of Kif18A KD cells expressing GFP-Kif18AFL and GFP-Kif18A^AVVVA^. **B-C**. Cells were treated with control (Control KD) or Kif18A siRNA (Kif18A KD) and transfected with the indicated GFP or GFP-Kif18A constructs. Cells were arrested in MG132 for 3 hours, then incubated with Hec1 phospho-Ser55 and total Hec1 antibodies and imaged. **B**. Plot of phospho-Ser55 Hec1 (pHec1) fluorescence intensity at kinetochores normalized to control phospho-Ser55 levels. **C**. Plot of mean normalized phospho-Ser55 Hec1 fluorescence for the indicated conditions. Error bars represent std dev. Means were compared using a student’s t-test. Data are from 3 independent experiments: n = 26 cells, 858 kinetochores (KTs), (Control KD, GFP only), n = 21 cells, 1188 KTs (Kif18A KD, GFP only), n = 13 cells, 705 KTs (Kif18A KD, GFP-Kif18AFL), n = 12 cells, 597 KTs (Kif18A KD, GFP-Kif18A^AVVVA^).

### Loss of Kif18A function leads to an increased number of unattached kinetochores

One of the consequences of higher Hec1 phosphorylation during prometaphase is that kinetochores are more likely detach from microtubules, even when they are properly bioriented [14, 20, 28]. Since mutating the PP1 binding site in the Kif18A C-terminus increased phospho-Hec1 levels at kinetochores in metaphase, we investigated whether these changes correlate with metaphase kinetochore-microtubule attachment defects. We used the presence of the SAC protein MAD1 at kinetochores, determined by colocalization with anti-centromere associated (ACA) antibodies, as a readout for unattached kinetochores [32]. Previous work indicates that loss of Kif18A leads to an increase in unattached kinetochores in asynchronously dividing HeLa cells [26]. We found that metaphase arrested Kif18A KD cells also display an increase in MAD1-positive kinetochores, indicating attachment errors persist to late metaphase in the absence of Kif18A (Figure S2A). The fraction of cells with at least one MAD1-positive kinetochore was similar in Kif18A KD cells expressing GFP-Kif18A^AVVVA^ (0.72 +/- 0.36, n = 42/58 cells) or GFP alone (0.8 +/- 0.4, n = 64/80 cells; Figure 3A-B). Control KD cells expressing GFP alone (0.23 +/- 0.12, n = 14/68 cells) had a lower fraction of cells with MAD1-positive kinetochores in comparison, as did cells expressing GFP-Kif18AFL (0.42 +/- 0.4, n = 22/50 cells; Figure 3A-B). Additionally, Kif18A KD cells expressing GFP-Kif18A^AVVVA^ have a similar number of MAD1-positive kinetochores per cell (3.9 +/- 4) as those expressing GFP (4.6 +/- 4) MAD1, while control KD cells expressing GFP (0.7 +/- 2) and Kif18A KD cells expressing GFP-Kif18AFL (1.6 +/- 3 per cell) both displayed significantly fewer MAD1-positive kinetochores per cell (Figure 3C). We found that cells with the highest levels of GFP-Kif18AFL expression had the fewest MAD1 positive kinetochores, but this trend was not true for GFP-Kif18A^AVVVA^ expressing cells (Figure S2B). Taken together, these data suggest that Kif18A-PP1 binding is necessary to maintain robust kinetochore-microtubule attachments and silence the SAC.

**Figure 3.**
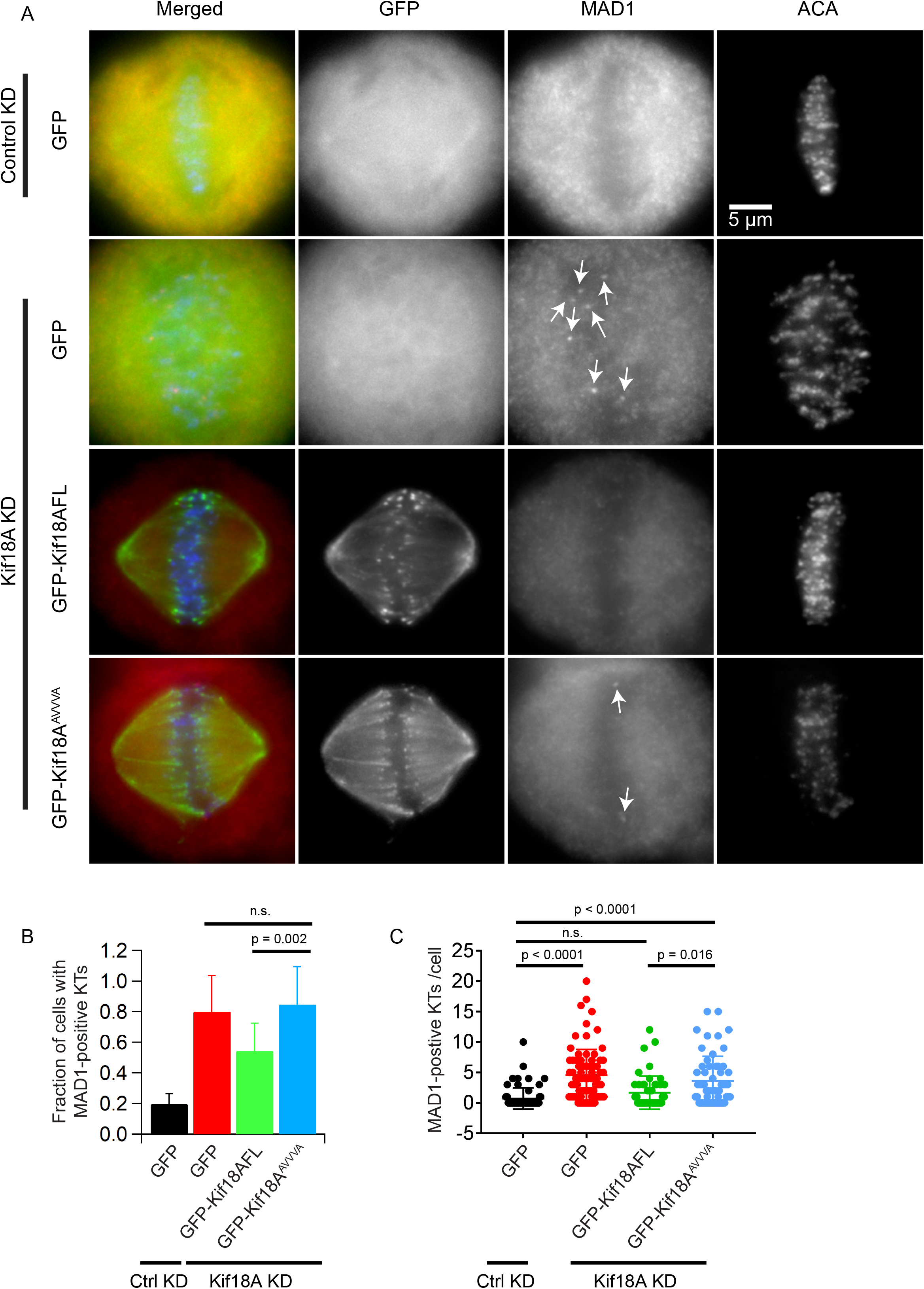
Metaphase cells expressing Kif18A^AVVVA^ display an increase in kinetochore-localized MAD1 on aligned kinetochores. **A**. Representative immunofluorescence images of Control KD or Kif18A KD HeLa cells expressing the indicated GFP constructs and immunofluorescently labeled for MAD1 and kinetochores (ACA). Arrows indicate MAD1-positive kinetochore. **B**. Bar graph displaying the fraction of cells with at least 1 MAD1-positive kinetochore (KT) for the indicated conditions. Data were compared using a chi-squared analysis. **C**. Scatter plot showing the number of MAD1-positive kinetochores per GFP-positive cell. Data were compared via a oneway ANOVA and Tukey’s multiple comparisons test. Data are from 3 independent experiments: n = 68 cells (Control KD, GFP only), n = 80 cells (Kif18A KD, GFP only), n = 50 cells (Kif18A KD, GFP-Kif18AFL), n = 58 cells (Kif18A KD, GFP-Kif18A^AVVVA^).

### Kif18A is capable of accumulating at K fiber plus-ends and aligning chromosomes independent of PP1 binding

We observed that metaphase arrested Kif18A KD cells expressing GFP displayed chromosome alignment defects similar to those previously reported in Kif18A loss of function cells, while Kif18A KD cells expressing GFP-Kif18A^AVVVA^ had comparatively well aligned chromosomes. To quantify chromosome alignment, we measured kinetochore distribution across the pole-to-pole axis of the spindle in Kif18A KD cells expressing GFP or GFP-Kif18A constructs [29, 33] (Figure 4A-B). All values were normalized to the average kinetochore distribution measured in GFP expressing cells treated with control siRNAs for comparison. We found that chromosome alignment in cells expressing either GFP-Kif18A^AVVVA^ or GFP-Kif18AFL was comparable to that seen in control cells (Figure 4C). In contrast, the kinetochore distribution in Kif18A KD cells expressing GFP alone was significantly wider than the distribution in Control KD cells (Figure 4C). These data indicate that Kif18A’s ability to align chromosomes, which depends on attenuation of microtubule dynamics, does not require PP1 binding. Consistent with this conclusion, we also find that GFP-Kif18A^AVVVA^ and GFP-Kif18A-FL accumulated with similar kinetics at the plus-ends of K fibers treated with the microtubule-stabilizing drug taxol, indicating that the plus-end directed motility and stable plus-end binding of the two motors is comparable (Figure S3). Stable microtubule plus-end accumulation is required for Kif18A’s function in suppressing microtubule dynamics, thus its role in chromosome alignment [33, 34]. Collectively, these data suggest that PP1-binding is not required for Kif18A’s motility or chromosome alignment functions during metaphase.

**Figure 4.**
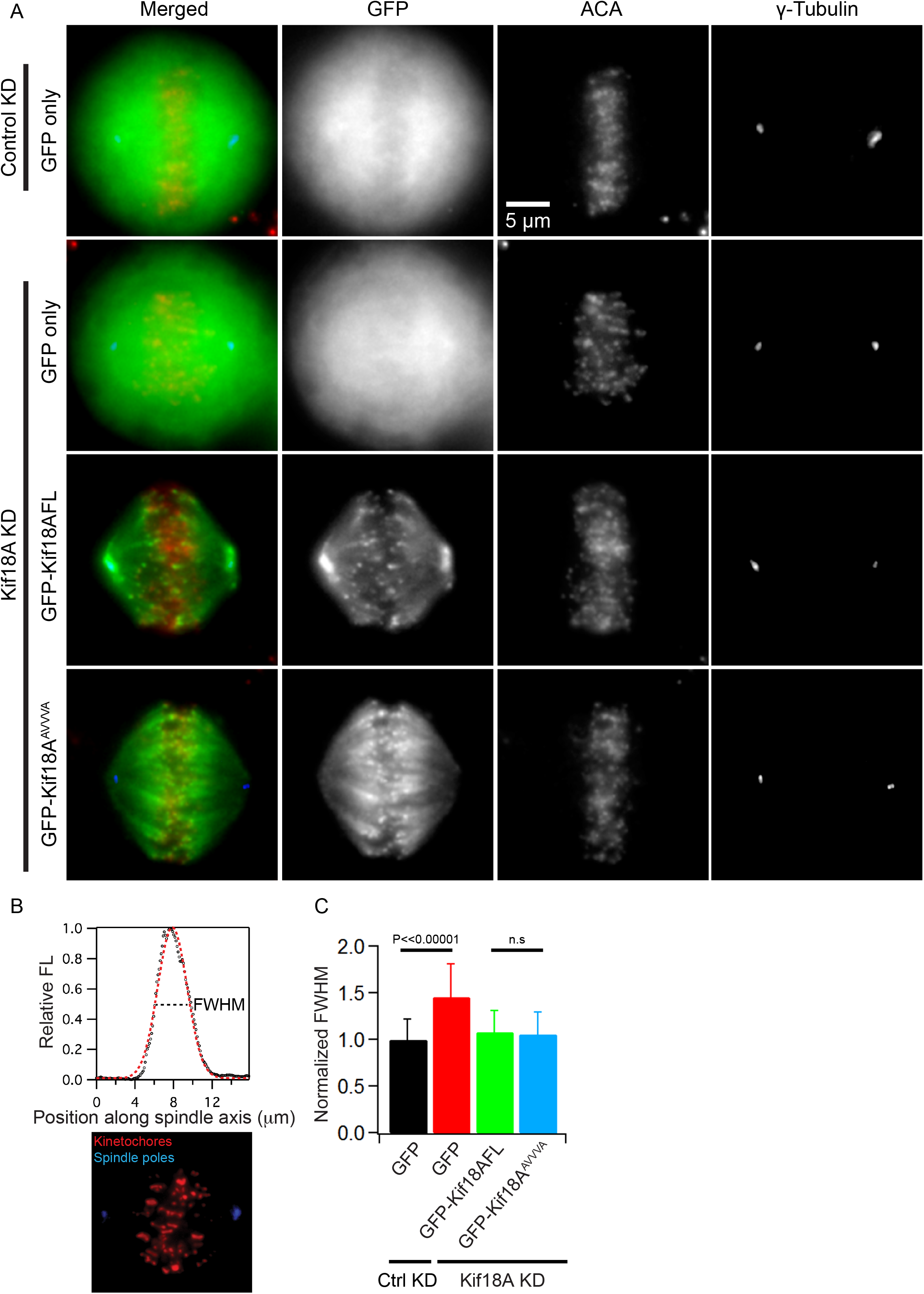
Kif18A is capable of aligning chromosomes independently of PP1 binding. A. Representative images of cells expressing GFP, GFP-Kif18AFL, or GFP-Kif18A^AVVVA^ variants immunoflourescently labeled for kinetochores (ACA) and centrosomes (γ-tubulin). Cells were treated with control (Control KD) or Kif18A siRNAs (Kif18A KD) and transfected with the indicated GFP constructs. **B**. Schematic of the method used to quantify kinetochore fluorescence distribution along the pole-to-pole axis of the spindle. Kinetochore fluorescence was fit to a Gaussian, and full width at half maximum (FWHM) was determined as a measure of chromosome alignment (dashed line in plot). **C**. A bar graph displaying normalized FWHM means for kinetochore fluorescence distributions from the indicated cell populations. Data were compared using a student’s t-test.

### Kif18A does not promote attachments by regulating chromosome positioning or kinetochore tension

It was previously reported that SAC proteins preferentially localized to unaligned kinetochores in Kif18A KD cells [26]. This raises the possibility that it is the unaligned chromosome population in these cells that has kinetochore microtubule attachment defects. Consistent with this idea, recent studies indicate that pole proximal kinetochores can be phosphorylated by Aurora A kinase, which has been shown to contribute to Hec1 phosphorylation [35, 36]. Therefore, we measured the location of MAD1 positive kinetochores relative to the center of the spindle in Kif18A KD cells expressing GFP-Kif18A constructs (Figure 5A). MAD-1-positive kinetochores were far from the midzone in Kif18A KD cells expressing GFP (2.59 +/- 1.54 μm) consistent with previous work [26] (Figure 5B). In contrast, MAD1-positive kinetochores in Kif18A KD cells expressing GFP-Kif18A^AVVVA^ were significantly closer to the midzone (1.36 μm +/- 96 μm) than those in Kif18A KD GFP cells and were positioned at comparable distances to those found in GFP-Kif18AFL (1.46 +/- 0.94 μm) and control KD GFP expressing cells (1.31 +/- 0.91 μm), which fully align their chromosomes (Figure 5B). These data indicate that chromosomes near the midzone require Kif18A-PP1 for attachment and that chromosome alignment is not sufficient for SAC inactivation.

**Figure 5.**
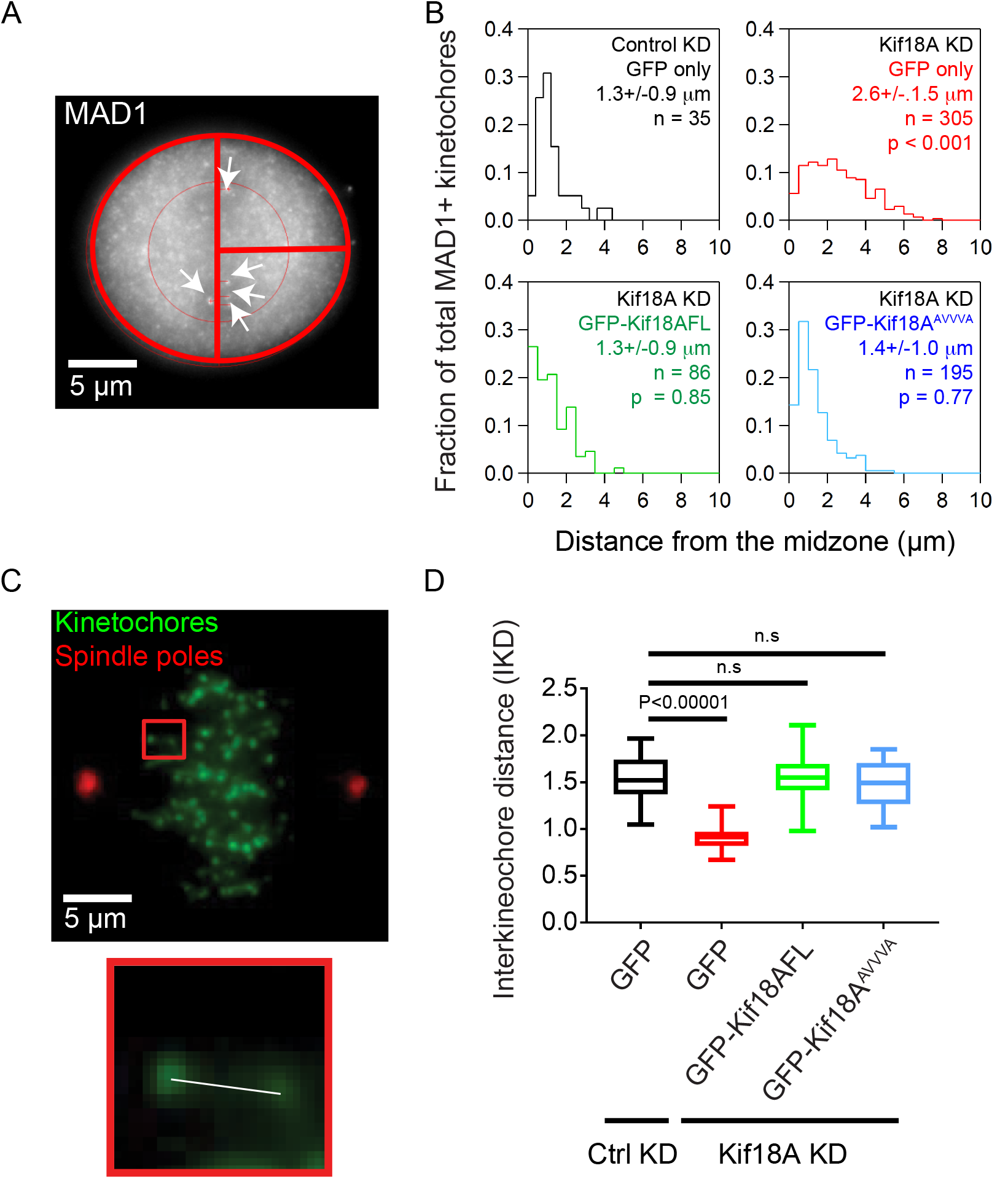
Kif18A does not promote attachments by regulating chromosome positioning or kinetochore tension. **A**. Representative image depicting the positions of MAD1-positive kinetochores (arrows) relative to the spindle midzone (vertical red line). **B**. Histograms displaying distances of MAD1 positive (MAD1+) kinetochores from the midzone for each condition listed. Mean +/- s.d. is indicated for each condition, n = number of kinetochores from >14 cells and 3 independent experiments. Data were compared using a one-way ANOVA with Dunnett’s multiple comparison test. Reported p-values represent comparison to Control KD + GFP condition. **C**. Mitotic cell immunofluorescently labeled for kinetochores (ACA) and spindle poles (γ-tubulin). Interkinetochore distance (IKD) was determined by measuring the distance between the centroids of sister kinetochores in the same focal plane (white line in inset). **D**. Box and whisker plot displaying the average IKD for the indicated experimental conditions. Data are from 3 independent experiments: n = 28 cells, 368 kTs (Control KD, GFP only), n = 55 cells, 531 kTs Kif18A KD, GFP only), n = 20 cells, 156 kTs (Kif18A KD, GFP-Kif18AFL), n = 21 cells, 176 kTs (Kif18A KD, GFP-Kif18A^AVVVA^). Data were compared using a student’s t-test.

Kif18A has also been implicated in regulating tension between kinetochores. Kif18A overexpression leads to hyperstretching between sister kinetochores, while depletion leads to a measurable decrease in interkinetochore distance (IKD) [4, 26]. Reduced kinetochore tension promotes Aurora B kinase-dependent phosphorylation of outer kinetochore substrates, such as Hec1 [37]. Thus, we measured the effect of GFP-Kif18A^AVVVA^ on IKD to determine if low tension could be contributing to the attachment defects observed in these cells. However, we found that the distances between sister kinetochores in Kif18A KD cells expressing GFP-Kif18A^AVVVA^ were comparable to those measured in control KD cells and Kif18A KD expressing GFP-Kif18AFL (Figure 5C-D). In contrast, Kif18A KD cells expressing GFP had significantly reduced IKDs compared to control KD cells (Figure 5D), consistent with previous reports [4, 26]. These results suggest that the attachment defects observed in GFP-Kif18A^AVVVA^ cells are not explained by reduced interkinetochore stretch.

### A low phosphorylation mimetic Hec1 mutant is sufficient to promote progression through mitosis in Kif18A KD cells

To determine if the SAC-dependent arrest observed in Kif18A KD cells is caused by increased Hec1 phosphorylation, we asked if a Hec1 mutant that mimics low phosphorylation could rescue mitotic progression. Specifically, we analyzed the effects of a previously characterized Hec1 mutant, called Hec1-1D, which carries alanine substitutions in eight of the predicted Aurora B phosphorylation sites and a single phosphomimetic aspartic acid substituted for Ser55 [15, 16]. Expression of GFP Hec1-1D in cells depleted of endogenous Hec1 permits normal chromosome movement but does not fully rescue chromosome alignment [16]. We co-depleted HeLa cells of Kif18A and Hec1 using previously validated siRNAs [4, 14], and subsequently transfected with plasmids encoding GFP-tagged wild type Hec1 (GFP-Hec1-WT), GFP-Hec1-1D, or GFP alone. The time from nuclear envelope breakdown to anaphase onset was measured using time-lapse DIC imaging in cells positive for GFP-Hec1 constructs (Figure 6A). The majority of Hec1 and Kif18A co-depleted cells expressing either GFP or GFP-Hec1-WT failed to divide during the time course of the movie or underwent apoptosis (Figure 6B). In contrast, the majority of GFP-Hec1-1D expressing cells completed division without chromosome alignment (Figure 6B). Thus, alleviating Hec1 phospho-regulation mitigates the mitotic arrest but not the chromosome alignment defects observed in Kif18A KD HeLa cells. These data suggest that a Kif18A-dependent increase in Hec1 affinity to microtubules silences the SAC and promotes the metaphase-to-anaphase transition independently of the motor’s known role in regulating microtubule dynamics.

**Figure 6.**
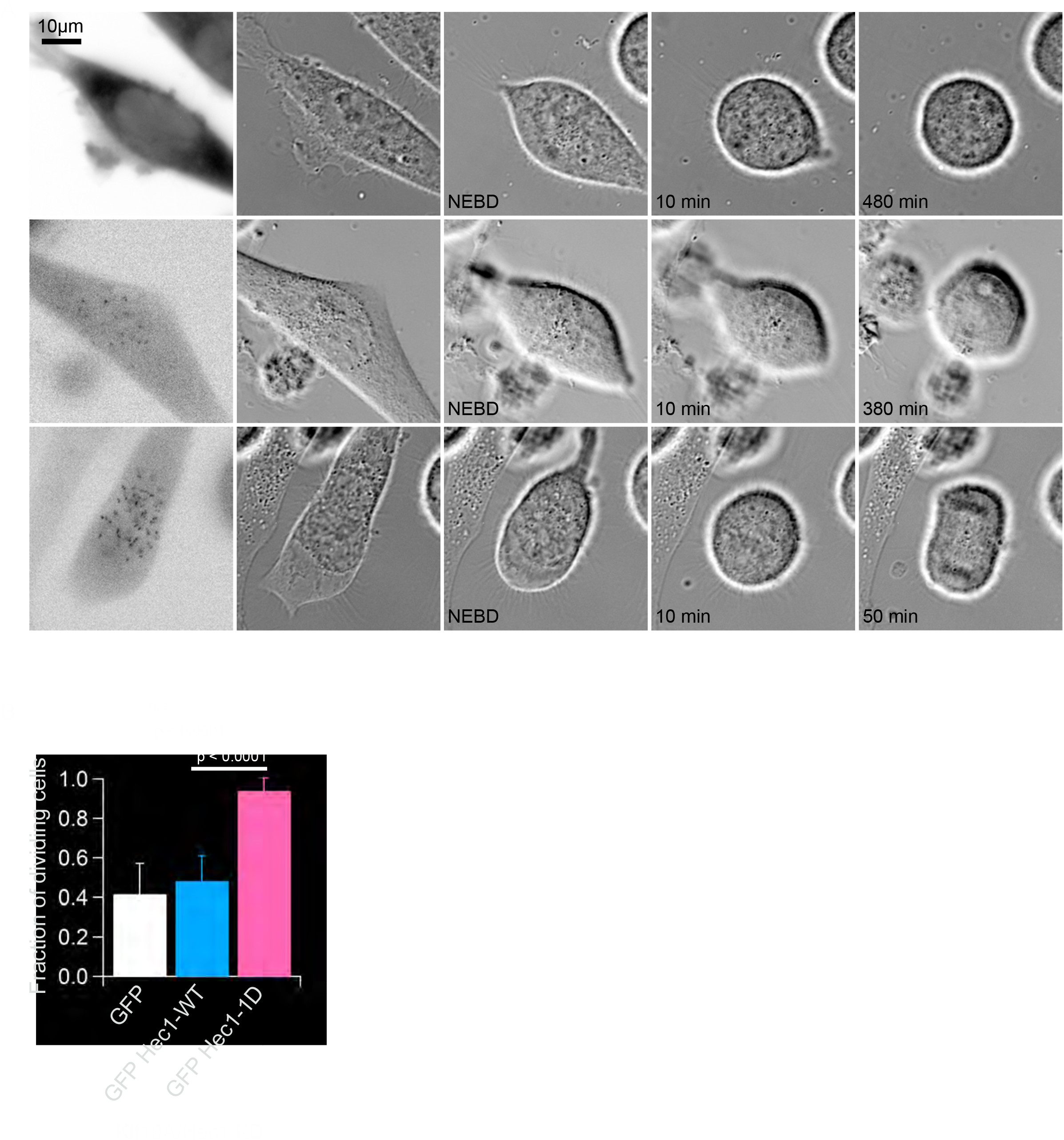
A Hec1 mutant that mimics low phosphorylation rescues the mitotic arrest caused by loss of Kif18A function. **A**. Still frames from time-lapse DIC imaging used to measure division in HeLa cells co-depleted of Hec1 and Kif18A and expressing GFP, GFP-Hec1-WT, or a GFP-tagged Hec1 mutant that mimics phosphorylation at a single site (GFP Hec1-1D). Mitotic division was defined as progress from nuclear envelope breakdown (NEBD) to anaphase onset. **B**. Bar graph representing the fraction of cells that divide in each condition. Data are from 3 independent experiments: n = 153 (GFP only), n = 79 (GFP Hec1-WT), and n = 70 (GFP Hec1-1D). Data were compared using chi-squared analyses.

## DISCUSSION

Biorientation and chromosome alignment during cell division ensure maintenance of genomic stability. Temporally regulated changes in the affinity between kinetochores and microtubules help to promote and stabilize bioriented attachments. As biorientation occurs, K fiber dynamics are also dampened to confine chromosome movements at the spindle midzone and form the metaphase plate. Our work indicates that Kif18A possesses two separable functions, one relying on its ability to suppress microtubule dynamics and the second on its direct interaction with PP1, that together couple the stabilization of bioriented attachments with the alignment of chromosomes at the spindle equator [22, 23].

Our data strongly support a model in which Kif18A promotes Hec1 dephosphorylation by recruiting PP1 to the plus-ends of K fibers (Figure 7). Our data indicate that the majority of kinetochores are attached to K fibers and devoid of SAC proteins during metaphase in the absence of Kif18A activity, suggesting that the motor is not necessary for initial end-on attachments during prometaphase (Figure 7A). This is also consistent with previous studies indicating that Kif18A’s motility to K fiber plus-ends, which is relatively slow (~75-100 nm/sec), is contingent on the presence of a stable kinetochore microtubule track (Figure 7B) [34, 38]. As kinetochores become attached, Kif18A accumulates at K fiber plus-ends, where it confines chromosome movements and promotes the complete dephosphorylation of Hec1. We propose that Kif18A’s dual functions provide positive feedback for K fiber attachment while the motor simultaneously dampens K fiber dynamics to align chromosomes during metaphase (Figure 7C-D). These combined activities facilitate temporal coordination between attachment and alignment.

**Figure 7.**
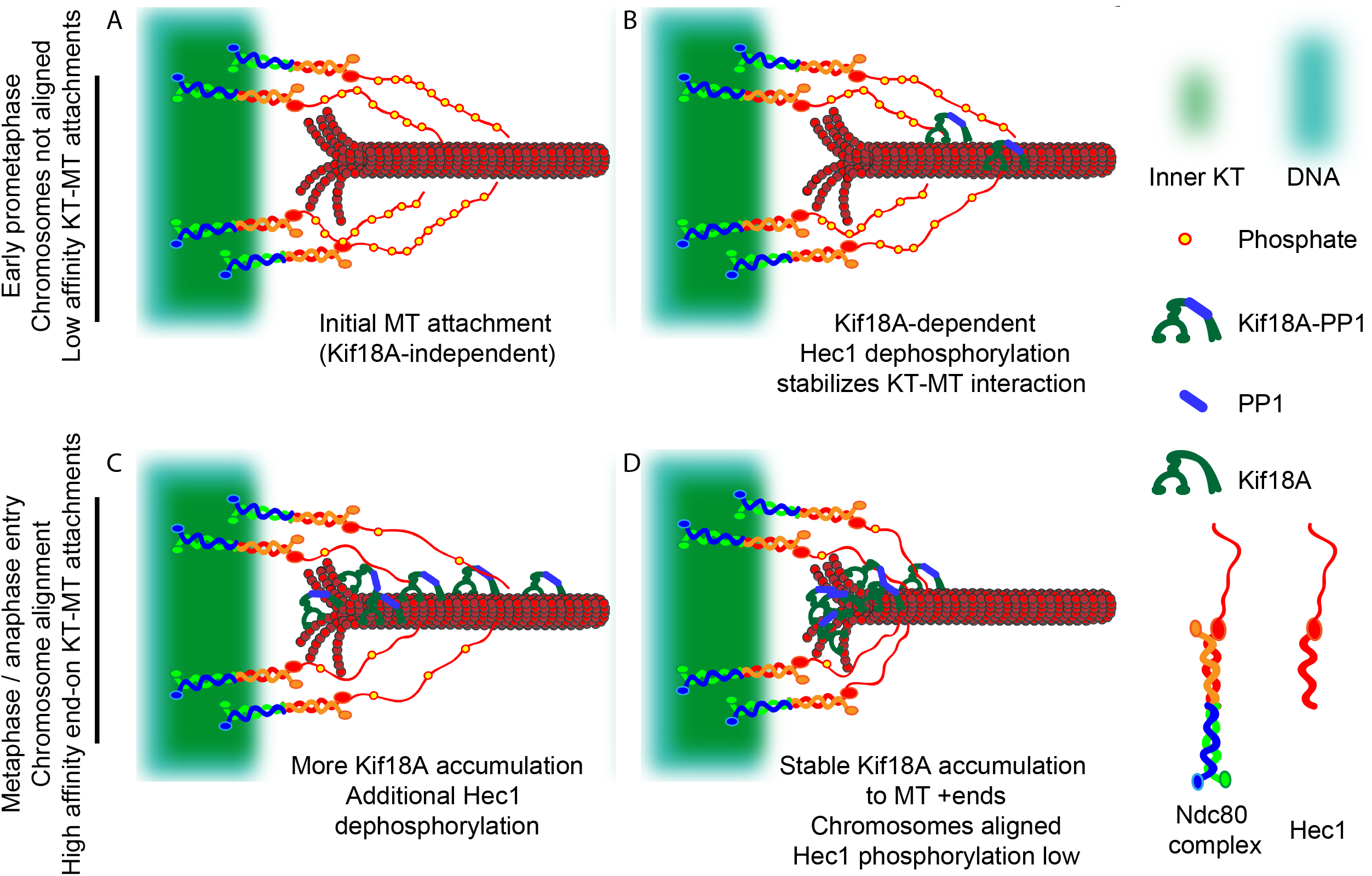
Model for Kif18A-dependent temporal coordination of chromosome alignment with stabilization of kinetochore microtubule attachments. **A**. During early prometaphase, microtubules (MTs) come in contact with the N-terminal tail of Hec1 in a Kif18A-independent manner. Phosphorylation of the Hec1 N-terminus is relatively high (yellow circles). **B**. Initial kinetochore microtubule (KT-MT) attachments (Kif18A-independent) provide stable tracks for Kif18A and Kif18A associated PP1 (Kif18A-PP1) to translocate to microtubule plus-ends. Kif18A-PP1 dependent Hec1 dephosphorylation reinforces KT-MT attachments. **C**. Kif18A-PP1 accumulation at KT-MT plus-ends dampens MT dynamics and confines chromosome movements to the spindle equator. In parallel, additional Kif18A-PP1 continues to dephosphorylate Hec1, providing positive feedback for KT-MT attachments. **D**. As K fibers mature, more Kif18A motors accumulate at K fiber plus-ends, promoting complete chromosome alignment and further enhancing stabilization of end-on attachments to silence the SAC and promote anaphase entry.

Recent findings show that Hec1 has different binding configurations on polymerizing and depolymerizing microtubules, and that Hec1 phosphorylation only affects microtubule association on polymerizing microtubules [39]. These data suggest that Hec1 dephosphorylation primarily increases the affinity of kinetochores for polymerizing K fibers. Kif18A’s behavior on growing and shortening microtubules is consistent with a role for the motor in specifically promoting the attachment of polymerizing K fibers. For example, Kif18A localizes asymmetrically on sister K fibers in mitotic cells [4]. Furthermore, purified Kif18A accumulates on growing microtubules but dissociates from shortening microtubules *in vitro* [34]. Taken together, these data imply that Kif18A may accumulate preferentially on growing K fibers, where it could promote Hec1 dephosphorylation. Switching to a phosphorylation-independent binding state during depolymerization circumvents the need for constant suppression of Aurora B activity near its outer kinetochore substrates for stabilizing and maintaining attachments.

Our data also address two alternative models for Kif18A’s role in promoting kinetochore microtubule attachment. We considered that Kif18A could indirectly promote Hec1 dephosphorylation by increasing tension at kinetochores, displacing Hec1 from the inner-centromere localized Aurora B kinase [37]. Consistent with this model, Kif18A accumulation at the plus-ends of K fibers enhances inter-kinetochore stretch [4, 26]. However, we found that cells expressing GFP-Kif18A^AVVVA^ displayed normal inter-kinetochore stretch but abnormally high Hec1 phosphorylation and MAD1 levels, suggesting Kif18A-dependent regulation of kinetochore tension does not explain its role in promoting Hec1 dephosphorylation or kinetochore microtubule attachment.

We also considered the possibility that Kif18A’s function in chromosome alignment excludes chromosomes from the spindle poles, preventing Aurora A kinase from phosphorylating Hec1. Pole proximal chromosomes are often improperly attached, and cells rely on Aurora A kinase activity to destabilize erroneous attachments [35, 36]. Some chromosome pairs in Kif18A depleted cells are significantly displaced from the metaphase plate. Thus, these misaligned pairs could be subject to regulation by Aurora A, explaining the observed increase in Hec1 phosphorylation. However, our data indicate that Kif18A KD cells expressing GFP-Kif18A^AVVVA^ have aligned chromosomes with increased phospho-Hec1 levels. Furthermore, primary embryonic fibroblasts isolated from *Kif18a* mutant mice divide with normal kinetics despite chromosome alignment defects [27]. These data collectively indicate that Kif18A’s chromosome alignment and attachment functions are independent of each other. While these alternative models are not mutually exclusive with Kif18A’s recruitment of PP1 to kinetochores, our data strongly support PP1-dependent dephosphorylation of Hec1 as the primary mechanism for Kif18A’s role in enhancing kinetochore-microtubule attachments.

Existing evidence suggests that a role for kinesin-PP1 coupling of chromosome alignment and attachment is conserved in eukaryotes. We have shown that murine primordial germ cells require Kif18A activity for kinetochore microtubule attachment, chromosome alignment, and completion of mitosis. However, while somatic cells from *Kif18a* mutant embryos also display chromosome alignment defects, they progress through mitosis with normal timing. These data are consistent with a role for Kif18A in coordinating chromosome alignment and attachment in at least some cell types during mammalian development and suggest that other mechanisms are able to compensate for the loss of Kif18A’s attachment function in others [18, 19, 40–42]. The fission yeast kinesin-8 motors Klp5/Klp6 are also required for chromosome alignment [42, 43] and interact with PP1 to silence the SAC [24]. The mechanism for checkpoint silencing in this case has not been identified, but Klp5/6 mutants are synthetically lethal with mutations in dam1 [44], suggesting a role for the heterodimeric Klp5/6 motor in kintetochore microtubule attachment. Furthermore, recent work from Suzuki *et al*. (accompanying paper) indicates that an interaction between Cin-8 (kinesin-5) and PP1 is required for kintetochore microtubule attachment in budding yeast. Interestingly, it is Cin-8, rather than the budding yeast kinesin-8 motor Kip3, that plays a more prominent role in regulating kinetochore microtubule dynamics and chromosome alignment in S. cerevisiae, suggesting Cin-8 may function to coordinate alignment and attachment [45].

In summary, our study indicates that the kinesin-8 motor Kif18A recruits PP1 and directly regulates microtubule dynamics to couple the attachment and alignment of mitotic chromosomes. Hec1 is a critical substrate of Kif18A associated PP1, and Kif18A-dependent dephosphorylation of Hec1 is required for cells to progress from metaphase into anaphase. In contrast, Kif18A-dependent chromosome alignment is not a prerequisite for satisfying the SAC. Thus, chromosome alignment and attachment are separable processes. Mechanisms that temporally coordinate these functions are likely necessary to prevent chromosome segregation errors.

## Experimental Procedures

### Plasmids and siRNAs

A Kif18A PP1 binding mutant (GFP-Kif18A^AVVVA^) was generated by mutating residues K612 and W616 via site-directed mutagenesis, resulting in K612A and W616A (forward sequence 5’-T C GAACATTT GGTAGAGAG GAAAGCAGT G GT AGTT GCGGCT GAC CAAACT GCC GA AC-3’) using siRNA-resistant GFP-Kif18AFL as a template, described previously in Stumpff *et al*, 2008. Cells were treated with previously validated control (Silencer Negative Control #1, Life Technologies), Kif18A (5’-GCUGGAUUUCAUAAAGUGG -3’, Life Technologies, [4, 29, 33], and Hec1 siRNAs (5-CCCUGGGUCGUGUCAGGAA-3’, Qiagen, [14].

### Cell culture and transfections

HeLa cells were cultured in 5% CO_2_ at 37°C in MEM-alpha (Life Technologies) with 10% fetal bovine serum (FBS, Gibco) and 1% antibiotics (Pen/Strep, Gibco). For siRNA transfections, 5.0 × 10^5^ cells were plated on 60mm^2^ dishes and grown overnight. 300 pmols of siRNA per 60mm^2^ dish (control and Kif18A) were incubated with Lipofectamine RNAiMAX transfection reagent (Thermo Scientific) in Opti-Mem reduced serum media as per manufacturer’s instructions, and added to cells for 8 hours. Cells were trypsinized, collected and pelleted. 5.0 × 10^5^ cells were electroporated with 3.5 μg of plasmid DNA (GFP only (pmax-GFP^TM^, Lonza, Germany), GFP-Kif18AFL, or GFP-Kif18A^AVVVA^, GFP-Hec1-WT, GFP-Hec1-1 D, or 2.0 μg of mCherry-PP1y, mCherry-CENP-B) using the Lonza 4D nucleofector system. Electroporated cells were seeded onto 12 mm^2^ glass coverslips (Electron Microscopy Sciences, Hatfield, PA) or Poly L-lysine treated glass bottom imaging dishes (MatTek) and allowed to incubate for approximately 18-24 hrs after transfection. Cells were treated with 20 μM MG132 for 3 hours to enrich for metaphase cells in fixed cell assays.

### Immunofluorescence

Cells were fixed on 12 mm^2^ glass coverslips (EMS) for 10 minutes in -20° C methanol and 1% paraformaldehyde on ice, and washed twice in TBS for 5 minutes. Coverslips were incubated with 20% boiled goat serum diluted in antibody-diluting buffer (Abdil, 1x TBS, 2% goat serum albumin, 0.2% sodium azide) for 1 hr at ambient temperature, then washed twice with TBS for 5 min. Coverslips were incubated with human anti-centromeric protein A (ACA) serum (Antibodies Incorporated, Davis, CA) at 2.5 μg/mL overnight at 4°C, mouse anti γ-tubulin at 1.0 μg/mL (Sigma-Aldrich, St. Louis, MO), mouse anti-α-tubulin at 1 μg/mL (Sigma-Aldrich), mouse anti-γ-tubulin at 1 μg/mL (Life technologies) at ambient temperature for 1 hr, mouse anti-MAD1 at 3.6 μg/mL at 4°C overnight. Cells used to detect total Hec1 and phospho-serine 55 Hec1 were pretreated with PHEM extraction buffer (60mM PIPES, 25mM HEPES, 10mM EGTA, 4mM MgCl_2_ (PHEM), 0.5% Triton-X, pH 6.9, 1x Halt protease and phosphatase inhibitor cocktail (Thermo Fisher) 100 μM Microcystin-LR (Sigma-Aldrich)) for 5 minutes at 37°C before fixing with 4% paraformaledehyde in 1x PHEM buffer for 15 minutes at ambient temperature. Cells were washed twice with 1x PHEM buffer for 5 min, then incubated at 4°C overnight with 3.44 μg/mL mouse anti-Hec1 (GeneTex) and 2.57 μg/mL rabbit anti-Hec1 Phospho-Ser55 (GeneTex). Coverslips were mounted onto glass slides with Prolong gold anti-fade mounting medium with 4’,6-Diamidine-2’-phenylindole dihydrochloride (Prolong Gold with DAPI, Life Sciences).

### Microscopy

Fixed and live cell imaging was performed using a Nikon Ti-E inverted microscope controlled by NIS Elements software (Nikon Instruments, Melville, NY) with APO 100x/1.49 numerical aperture (NA) and Plan APO 60x/1.42 NA oil immersion objectives (Nikon Instruments), Spectra-X light engine (Lumencore, Beaverton, OR), and Clara cooled charge-coupled device (CCD) camera or iXon X3 DU-897 EMCCD camera (Andor, South Windsor, CT). Some fixed cell imaging was also performed using a Nikon TE2000-E2 inverted microscope controlled by NIS elements software (Nikon Instruments) with a Plan Apo 60X/1.42 NA oil immersion objective, EXFO X-CITE 120 illuminator, Uniblitz shutters, and a Photometrics Cool-SNAP HQ2 14-bit camera.

### Live imaging of mitotic division

Cells co-depleted of Kif18A and Hec1 were electroporated with GFP-Hec1-WT or GFP-Hec1-1D plasmid DNA (kind gifts from Dr. Jennifer DeLuca, Zaytsev 2014) 8 hrs after siRNA incubation, then plated onto 35 mm^2^ glass bottom filming dishes (MatTek) and imaged approximately 30 hours post-electroporation. Cell culture media was changed to CO_2_-independent media containing 10% FBS and 1% antibiotics (Pen/Strep) prior to imaging. DIC images were taken every 2 minutes, with GFP images taken once every 40 frames concurrently to minimize phototoxicity. Only cells that were positive for GFP-Hec1 constructs were analyzed. For analyses of dividing cells, only cells that entered mitosis during the movie and displayed one of the following behaviors were included: (1) completed division during the movie, (2) underwent apoptosis (counted as did not divide), or (3) were in mitosis for at least two hours and were still in mitosis at the end of the movie (counted as did not divide).

### Chromosome alignment analyses

After fixation, single focal plane images were taken of mitotic cells with both spindle poles in the same plane of focus. The images were rotated to ensure that the spindle pole axis was horizontal, and a line was drawn between the two poles. The Plot Profile command in ImageJ was used to measure the distribution of ACA-labeled kinetochore fluorescence within a region of interest (ROI) defined by the length of the spindle with a set height of 17.5 μm (Stumpff 2012, Kim 2014). Plots of normalized average ACA fluorescence in each pixel column as a function of distance along the normalized spindle pole axis were analyzed by Gaussian fits using Igor Pro (Wavemetrics). The full width at half maximum intensity for the Gaussian fit was used as a metric for chromosome alignment in that cell.

To measure the position of MAD1-positive kinetochores relative to the spindle midzone, an ellipse was drawn to encompass the mitotic cell. Major and minor axes were determined by the ellipse draw command in NIS elements, with major axes drawn parallel to the spindle pole axis. The midzone was determined as the point that bisects the major axis. Distances were measured from the midzone to MAD1-positive kinetochores with fluorescence intensity greater than 1.5 times the background fluorescence.

### Total Hec1 and phospho-Hec1 Ser55 fluorescence quantification

A maximum intensity projection of 5 optical sections taken at 200 nm intervals was used to quantify total Hec1 and phospho-Hec1 Ser55 fluorescence. Hec1 foci were used to define regions of interest (ROIs) at individual kinetochores where both total and phospho-Hec1 Ser55 fluorescence were measured. Only ROIs that did not overlap with a neighboring kinetochore were analyzed. All fluorescence measurements were normalized to the mean control fluorescence level in each channel.

### Motor accumulation measurements

Kif18A-depleted cells co-transfected with GFP rescue constructs (GFP, GFP-Kif18AFL, or GFP-Kif18A^AVVVA^) and mCherry-CENPB were plated on 35 mm^2^ glass bottom filming dishes and imaged approximately 30 hours post-transfection. Cell culture media was changed to CO_2_-independent media containing 10% FBS and 1% antibiotics (Pen/Strep) prior to imaging. To determine motor accumulation after taxol addition, an equal volume of 20 μM taxol diluted in CO_2_-independent media was added to each dish to achieve a final concentration of 10 μM taxol. Four frames were collected at 5 sec intervals prior to taxol addition. Following taxol addition, images were taken at 5 sec intervals for 5 minutes. To quantify the relative rate of motor accumulation to K fiber plus-ends, GFP fluorescence was fit to a Gaussian across the spindle pole axis at every frame of the movie. The full width at half maximum (FWHM) of GFP fluorescence was measured and plotted as a function of time to obtain a relative rate of accumulation, reported as a rate of decreasing FWHM over time.

## Acknowledgements

Support for this work comes from NIH GM121491 (to JS), Susan G Komen grant CCR16377648 (to JS), and an Institutional Research Grant 14-196-01 from the American Cancer Society. The authors wish to thank Cindy Fonseca for technical assistance, Guy Kennedy for microscope support, and Dr. Jennifer DeLuca for reagents. We also thank Dr. Laura Reinholdt, Dr. Heidi Malaby, Alex Thompson, Leslie Sepaniac, Cindy Fonseca, Dr. Jamie Stern, and Victoria DeVault for insightful comments and suggestions.

## Author Contributions

Conceptualization, HK and JS; Methodology, HK and JS; Investigation, HK and JS; Resources, HK and JS; Writing-Original Draft, HK and JS; Writing-Review & Editing, HK and JS; Visualization, HK and JS; Supervision, JS.

---

**Movie S1. Accumulation of GFP-Kif18AFL at kinetochore microtubule plus-ends following taxol addition**. Images were collected at 5 sec intervals and are played back at 25 frames per second.

**Movie S2. Accumulation of GFP-Kif18A^AVVVA^ at kinetochore microtubule plus-ends following taxol addition**. Images were collected at 5 sec intervals and are played back at 25 frames per second.

